# The D614G Mutation Enhances the Lysosomal Trafficking of SARS-CoV-2 Spike

**DOI:** 10.1101/2020.12.08.417022

**Authors:** Chenxu Guo, Shang-Jui Tsai, Yiwei Ai, Maggie Li, Andrew Pekosz, Andrea Cox, Nadia Atai, Stephen J. Gould

**Author notes:** Correspondence: Name: Stephen J. Gould Ph.D, Address: 725 North Wolfe Street, Baltimore, MD 21205.

## Abstract

The spike D614G mutation increases SARS-CoV-2 infectivity, viral load, and transmission but the molecular mechanism underlying these effects remains unclear. We report here that spike is trafficked to lysosomes and that the D614G mutation enhances the lysosomal sorting of spike and the lysosomal accumulation of spike-positive punctae in SARS-CoV-2-infected cells. Spike trafficking to lysosomes is an endocytosis-independent, V-ATPase-dependent process, and spike-containing lysosomes drive lysosome clustering but display poor lysotracker labeling and reduced uptake of endocytosed materials. These results are consistent with a lysosomal pathway of coronavirus biogenesis and raise the possibility that a common mechanism may underly the D614G mutation’s effects on spike protein trafficking in infected cells and the accelerated entry of SARS-CoV-2 into uninfected cells.

## Introduction

COVID-19 (coronavirus infectious disease 2019) is caused by infection with the enveloped virus SARS-CoV-2, a member of the betacoronavirus family (Coronaviridae Study Group of the International Committee on Taxonomy of, 2020; Zhou et al., 2020). In just one year, SARS-CoV-2 has infected >65 million people, killed >1.5 million people, and caused extensive morbidity among survivors. Of all the RNA viruses, coronavirus have the largest genomes, encoding more than two dozen proteins (Wang et al., 2020). However, leading vaccine candidates elicit immunity to just a single protein, spike (Poland et al., 2020), which mediates the first two steps in the replication cycle: cell binding and fusion of viral and cellular membranes.

SARS-CoV-2 entry leads to translation of its viral genomic RNA (gRNA) into the orf1a and orf1a/b polyproteins (V’Kovski et al., 2020). These are then processed to generate 16 non-structural proteins (nsps) that reprogram the cell for viral replication and drive the synthesis of subgenomic viral mRNAs. These encode the four major structural proteins of mature virions (nucleocapsid (N), spike (S), membrane (M), and envelope (E)) and nine additional ORFs (3a, 3b, 6, 7a, 7b, 8, 9b, 9c, & 10) that together drive virus particle assembly and release. The spike, membrane and envelope proteins are sufficient to drive the formation of virus-like-particles and are thought to drive virion budding as well. This occurs following the synthesis of these proteins in the endoplasmic reticulum (ER) and their vesicle-mediated transport to the ER-Golgi intermediate compartment (ERGIC) (Ruch and Machamer, 2012; Ujike and Taguchi, 2015). As for how fully-formed virus particles (Yao et al., 2020) are released from infected cells, the prevailing model has been that they are released by the biosynthetic secretory pathway (Ujike and Taguchi, 2015). However, Altan-Bonnet and colleagues recently demonstrated that egress of mouse hepatitis virus (MHV, another betacoronavirus) does not require the biosynthetic secretory pathway, and instead uses an Arl8-dependent, lysosomal pathway of egress (Ghosh et al., 2020).

Shortly after its entry into the human population, a variant strain of SARS-CoV-2 arose that displays multiple hallmarks of increased viral fitness, including a higher rate of transmission, elevated viral load *in vivo*, and enhanced infectivity *in vitro* (Hou et al., 2020; Korber et al., 2020; Lorenzo-Redondo et al., 2020) This variant contains a mutation in the spike gene, D614G, that replaces the aspartate at position 614 with a glycine. This mutation lies N-terminal to the polybasic site cleavage site (682RRAR685), a cleavage event that is essential to virus replication and generates the non-covalently associated N-terminal S1 and C-terminal S2 proteins (Hoffmann et al., 2020a; Hoffmann et al., 2020b). The spike S1 domain binds SARS-CoV-2 receptors, primarily angiotensin converting enzyme-2 (ACE2) (Hoffmann et al., 2020b; Matheson and Lehner, 2020; Zhou et al., 2020)(Wrapp et al., 2020) but also neuropilin-1 (Cantuti-Castelvetri et al., 2020; Daly et al., 2020), and yet, there is as yet no evidence that the D614G mutation alters the affinity of S1 for its receptors or for target cells. In fact, ultrastructural and biochemical analyses of matched D614 and G614 SARS-CoV-2 viruses show no differences in their appearance in electron micrographs, and also no difference in levels of virion-associated spike proteins (Hou et al., 2020).

Here we extend our understanding of the spike D614G mutation by demonstrating that it shifts spike protein trafficking towards the lysosome and away from organelles of the biosynthetic secretory pathway. This shift is reflected in the localization of SARS-CoV-2 spike protein when expressed on its own, outside the context of an infected cell, and also in the context of SARS-CoV-2 infected cells. These and other results raise the possibility that that a single, lysosome-related mechanism may explain the pleiotropic effects of the D614G mutation on spike protein biogenesis, accelerated virus entry, and enhanced infectivity.

## Results

### Validating IFM-based SARS-CoV-2 serology tests

Fluorescence microscopy offers many advantages as a serology test platform, including high sensitivity, broad dynamic range, use of proteins in their native conformation and biological context, ability to incorporate in-sample controls that minimize false positive and false negative results, and the potential to interrogate multiple antibody responses in a single sample. As a first step towards the development of such an assay for SARS-CoV-2 antibodies, we generated cell lines designed to express the SARS-CoV-2 nucleocapsid, spike, or membrane proteins in response to doxycycline. These Htet1/N, Htet1/S**, and Htet1/M cell lines were subjected to immunoblot analysis, which confirmed that each cell line expressed its cognate SARS-CoV-2 protein in response to doxycycline (***supplemental figure S1***). It should be noted that S** corresponds to full-length spike that carries a pair of trimer-stabilizing proline mutations (986KV987 to 986PP987) and a quartet of substitutions at the S1/S2 protease cleavage site (682RRAR685 to 682GSAG685), a form of spike that displays the prefusion conformation (Wrapp et al., 2020) and may therefore be particularly effective at capturing neutralizing anti-spike antibodies.

We next seeded Htet1/N, Htet1/S**, and Htet1/M cells into 96-well glass-bottom tissue culture plates along with ∼10% Htet1 cells (to serve as an internal negative control), incubated the cells overnight in doxycycline-containing media, and then processed these cells for immunofluorescence microscopy. In this format, the fixed adherent cells serve as a solid support for capturing anti-SARS-CoV-2 antibodies from patient plasmas, the presence of human antibodies bound to the viral protein is detected using fluorescently-tagged donkey antibodies specific to human immunoglobulins (IgG, IgM, and IA), mCherry identifies the viral protein-expressing cells, and DAPI stains the nucleus. When these cells were tested using 40 control plasmas (all collected prior to the COVID-19 pandemic), we failed to detect any specific reaction to the N, S** or M proteins (***Fig. 1, top row***; ***supplemental figures S2-S5***).

**Figure 1.**
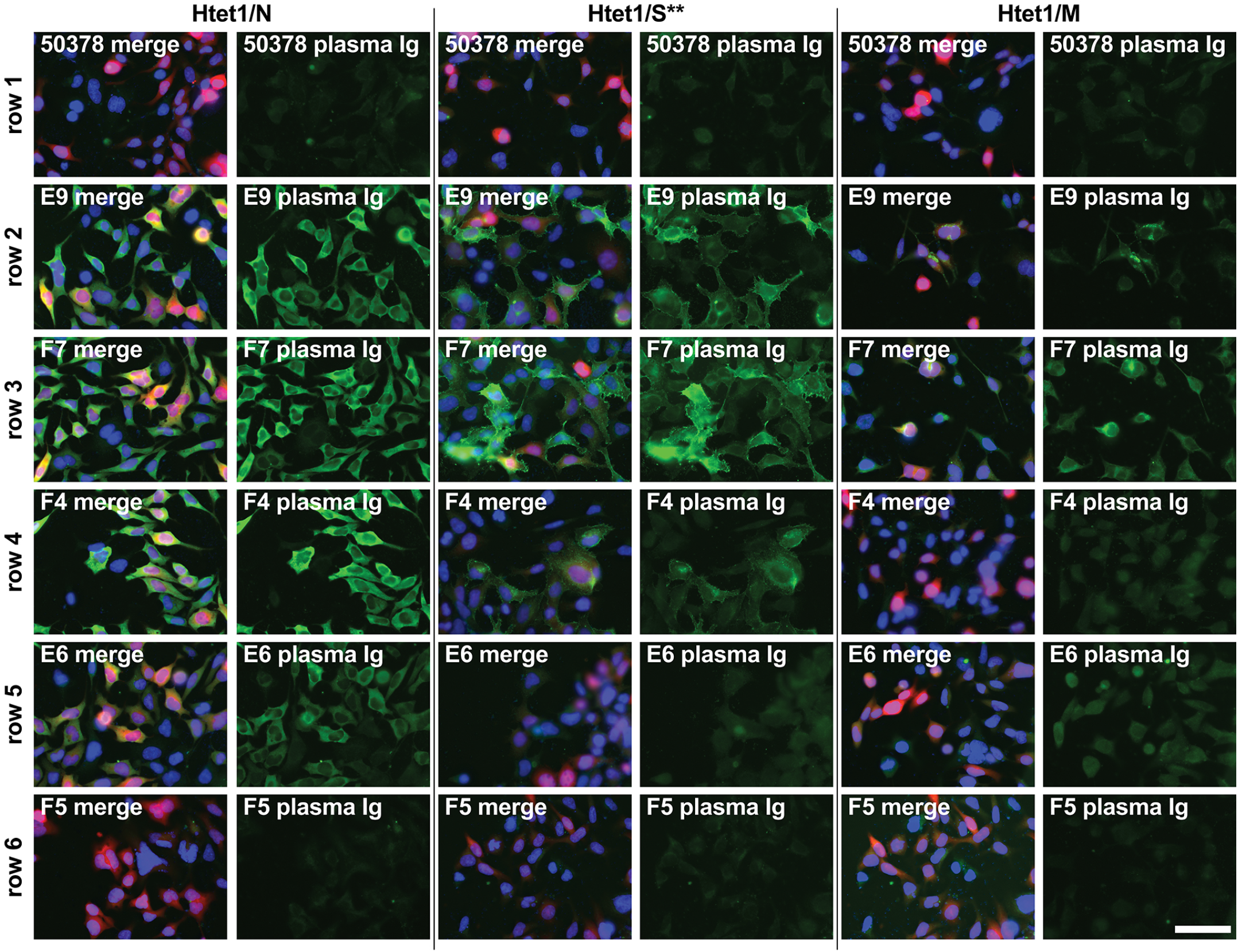
Microscopy-based serology. Fluorescence micrographs of (left two columns) Htet1/N, (middle two columns) Htet1/S**, and (right two columns) Htet1/M cells that had been processed for immunofluorescence using human plasmas from (row 1) a control patient and (rows 2-6) 5 different COVID-19 patients. Bound antibodies were detected using Alexafluor 488-conjugated anti-human Ig antibodies, DAPI was used to stain the nucleus, and mCherry was expressed from the same transposon as the viral protein. The left column in each pair shows the merge of all three images while the right column of each pair shows the staining observed for the patient plasma antibodies. Bar, 75 μm.

In contrast, plasmas from hospitalized, PCR-confirmed, COVID-19 patients (collected on day 0 of admittance into Johns Hopkins Hospital during April 2020) displayed a variety of reactivities towards the N, S**, and M proteins. Of the 30 COVID-19 patient plasmas that were tested, 23 (∼3/4) contained antibodies that bound to N-expressing cells, 20 (∼2/3) contained antibodies that bound to S**-expressing cells, and 13 (∼1/2) scored positive for the presence of anti-M antibodies (***Fig. 1, rows 2-6***; ***supplemental figures S6, S7***). Moreover, all plasmas with anti-M antibodies also contained anti-N and anti-S antibodies (***Table 1***). These results establish the validity of microscopy-based serology testing, indicate that the levels of anti-M antibodies may be a more reliable indicator of a deep and broad antibody responses to SARS-CoV-2 infection, and are generally consistent with prior studies of COVID-19 patient antibody responses (Grzelak et al., 2020; Whitman et al., 2020),

**Table 1.** Microscopy-based serology assay results for 30 COVID-19 plasmas.

| Plasma | α-N Ab | α-S Ab | α-M Ab |
| --- | --- | --- | --- |
| E5 | positive | positive | negative |
| E6 | positive | positive | negative |
| E7 | positive | positive | negative |
| E8 | positive | positive | positive |
| E9 | positive | positive | positive |
| E10 | positive | positive | negative |
| E11 | positive | positive | positive |
| E12 | positive | positive | positive |
| F1 | positive | positive | positive |
| F2 | positive | positive | negative |
| F3 | positive | positive | positive |
| F4 | positive | positive | negative |
| F5 | negative | negative | negative |
| F7 | positive | positive | positive |
| F8 | negative | negative | negative |
| F9 | positive | negative | negative |
| F10 | negative | negative | negative |
| F11 | negative | negative | negative |
| F12 | positive | positive | positive |
| G1 | positive | positive | positive |
| G2 | positive | positive | positive |
| G3 | negative | negative | negative |
| G4 | positive | positive | negative |
| G5 | positive | negative | negative |
| G6 | negative | negative | negative |
| G7 | positive | negative | negative |
| G8 | negative | negative | negative |
| G9 | positive | positive | positive |
| G10 | positive | positive | positive |
| plasma 5 | positive | positive | positive |

### Patient antibodies reveal an antigenically distinct subpopulation of spike

The SARS-CoV-2 spike protein is known to oscillate between multiple conformational states (Ke et al., 2020) and the anti-spike antibody responses of COVID-19 patients may allow us to detect functionally significant fluctuations in spike protein confirmation within its native context of the human cell. Given that the 986KV987 to 986PP987 and 682RRAR685 to 682GSAG685 substitutions present in the S** protein are designed to limit such conformational fluctuations (Wrapp et al., 2020), we developed cell lines that express functional forms of spike. Htet1/S^W1^ was engineered to express the spike protein encoded by the Wuhan-1 isolate of SARS-CoV-2 (Zhou et al., 2020) and was probed with plasmas with a small subset of COVID-19 patients (i.e. E12, E9, 5, and G4). Some of these plasmas (plasmas E12 and E9) generated an antibody staining pattern against the S^W1^ protein that conformed to the expected distribution, namely strong staining at the plasma membrane and in some cells in the Golgi, shown here by the occasional co-localization with GM130 (***Fig. 3, left column***). However, staining with other plasmas (5, G4) allowed the detection of yet another subpopulation of spike proteins within large, non-Golgi, intracellular structures.

**Figure 2.**
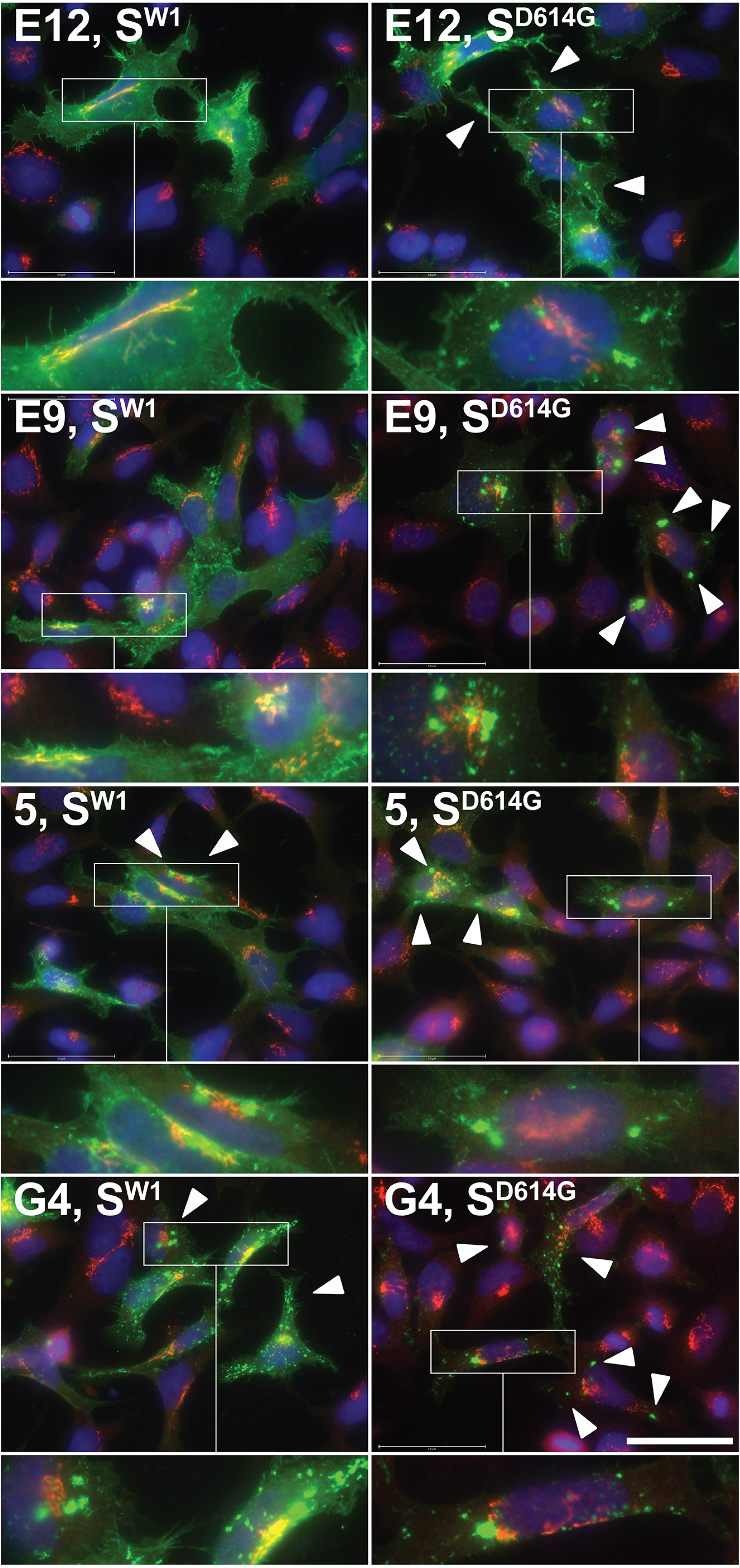
Human plasmas reveal differential trafficking of antigenically distinct forms of spike. Fluorescence micrographs of (left column) Htet1/S^W1^ cells and (right column) Htet1/S^D614G^ stained with (green) plasmas from COVID-19 patients E12, E9, 5, and G4, as well as with (red) antibodies specific for the Golgi marker GM130, and (blue) DAPI. White arrowheads point to large, spike-containing intracellular compartments that lack GM130. Insets (∼2.5-fold higher magnification) show greater detail in areas of particular interest. Bar, 50 μm.

**Figure 3.**
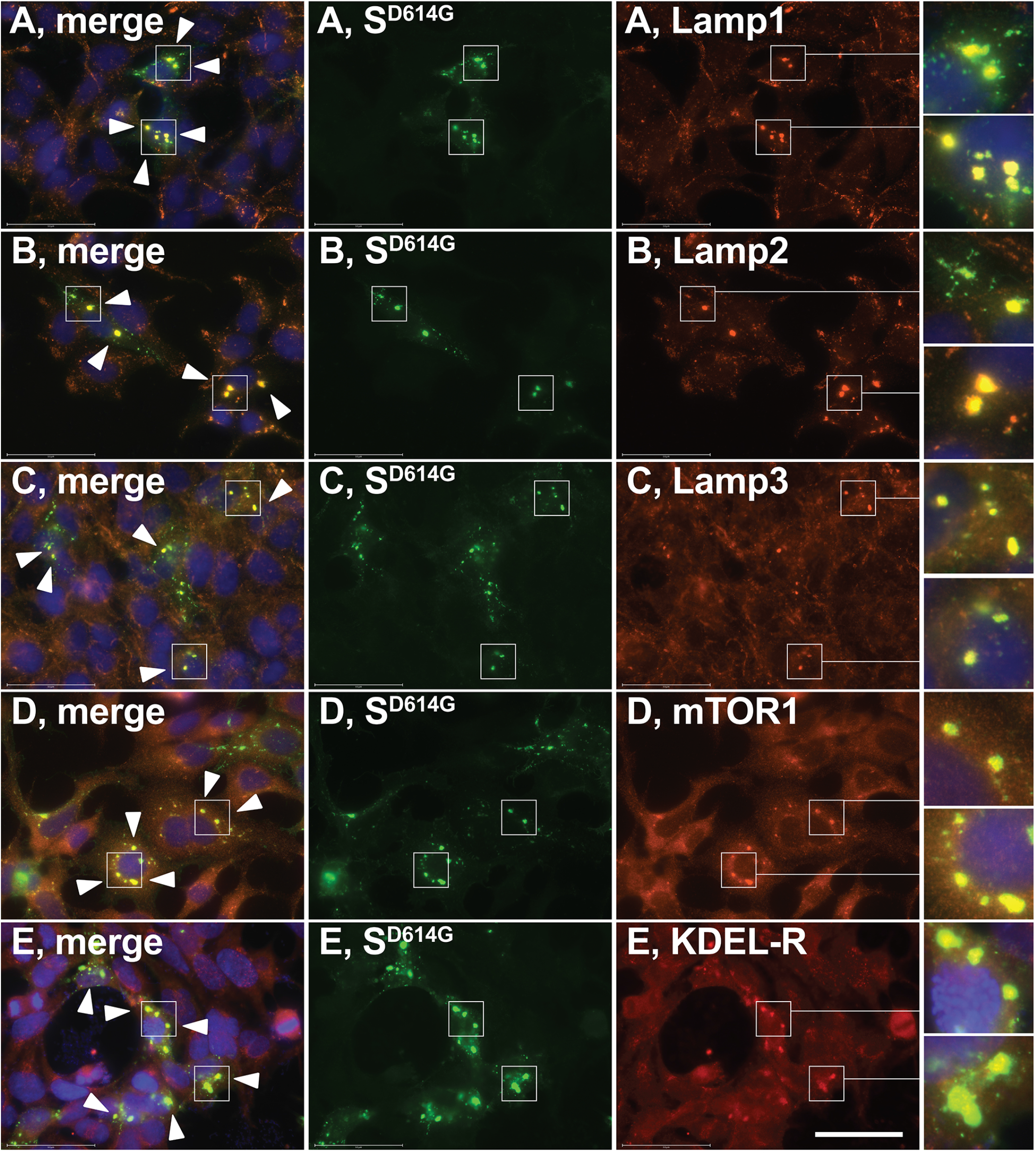
SARS-CoV-2 spike is trafficked to lysosomes. Fluorescence micrographs of Htet1/S^D614G^ cells induced with doxycycline and processed for immunofluorescence microscopy using (blue) DAPI, (green) plasma from COVID-19 patient G4, and (red) antibodies specific for for (A) Lamp1, (B) Lamp2, (C) Lamp3/CD63, (D) mTOR1, and (E) KDEL receptor (KDEL-R). White arrowheads denote the positions of S^D614G^-containing lysosome-related compartments. Insets (∼3-fold higher magnification) show greater detail in areas of particular interest. Bar, 50 μm.

These results indicate that spike expression in human cells leads to the creation of antigenically distinct subpopulations of spike that display different subcellular distributions, and furthermore, that COVID-19 patients vary significantly in their development of antibodies to these different conformations. In light of these findings, we tested whether plasma antibodies might shed light on the effects of the spike D614G mutation, which is known to enhance viral fitness through an increase in SARS-CoV-2 infectivity and transmission (Hou et al., 2020; Korber et al., 2020; Lorenzo-Redondo et al., 2020). Towards this end, we generated the Htet1/S^D614G^ cell line and interrogated S^D614G^-expressing cells using the same four plasmas (***Fig. 3, right column***). Although this form of spike differs from S^W1^ by just a single amino acid, all four plasmas revealed enhanced staining of spike in these large, non-Golgi intracellular compartments, raising the possibility that the D614G mutation either enhances the trafficking of spike to these compartments or enhances a conformational shift that allows its detection by the antibodies in these plasmas.

### SARS-CoV-2 spike is trafficked to lysosomes

Given the central role of spike in SARS-CoV-2 biology, the pronounced impact of the D614G mutation on the COVID-19 pandemic, and the fact that all leading vaccine candidates are based solely on spike, we sought to determine the identity of these large, spike-containing structures. This was done by inducing Htet1/S^D614G^ cells to express spike and then processing the cells for immunofluorescence microscopy using plasma G4 and antibodies directed against marker proteins of various subcellular compartments. These experiments showed no labeling of these structures with markers of the ER (calnexin and BiP), the ERGIC (ERGIC53 and ERGIC3,) the Golgi (GM130), endosomes (EEA1) or the plasma membrane (CD810 (***supplemental figure S8***). However, these structures did label with antibodies specific for lysosomal membrane markers Lamp1 and Lamp2 (***Fig. 3A, B***) and the lysosome-associated proteins Lamp3 (CD63) and mTOR (***Fig. 3C, D***). It has recently been shown that MHV-infected cells mislocalize the KDEL receptor (KDELR) from the ER and Golgi to the lysosome as part of virus-mediated reprogramming of host secretory pathways (Ghosh et al., 2020), and we observed that spike expression induced a redistribution of the KDEL receptor to the lysosome in a portion of spike-expressing cells (***Fig. 3E***).

These experiments revealed that lysosomes of spike-expressing cells accumulated in large clusters, a phenotype that was relatively uncommon in non-expressing cells. This appearance is likely due to lysosome clustering rather than lysosome fusion, and to test this hypothesis we examined S^D614G^ and Lamp2 distibution following a brief exposure to vacuolin-1, a PIKfyve inhibitor known to induce lysosome swelling (Huynh and Andrews, 2005; Sano et al., 2016). As expected, cells treated with vacuolin-1 contained swollen lysosomes several micrometers in diameter (***Fig. 4A-C***). Importantly, the membranes of these swollen lysosomes were labeled for both S^D614G^ and Lamp2, confirming that these proteins were indeed co-localized in lysosome membranes. Moreover, the sizes and numbers of these lysosomes appeared to be similar in expressing and non-expressing cells, indicating that spike expression induces lysosome clustering rather than lysosome fusion.

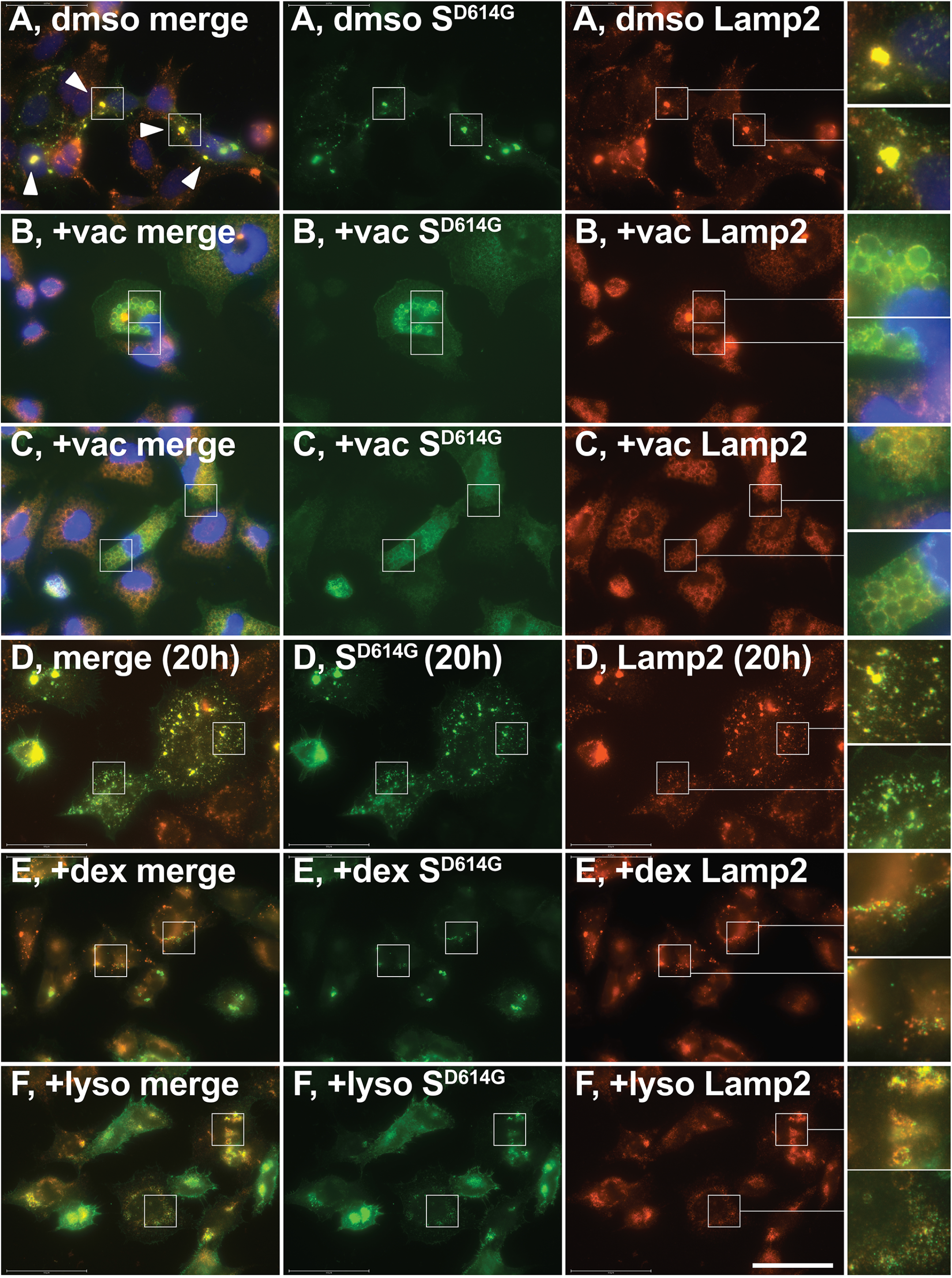

### Spike expression disrupts lysosome function

Lysosomes are the terminal destination for materials brought into the cell by fluid-phase endocytosis (Braulke and Bonifacino, 2009), a process that can be monitored by incubating cells with fluorescently-labeled, high molecular weight dextrans (Ohkuma, 1989). To determine whether spike-containing lysosomes retained this core function of lysosomes, we incubated Htet1/S^D614G^ cells with Alexa Fluor-647-labeled dextran (∼10 kDa), followed by a 3-hour chase in label-free media to allow endocytosed material to reach the lysosome. Furthermore, we performed these experiments on cells that had only been induced to express spike for 20 hours rather than several days, to increase the probability that we visualize cells at an earlier stage of lysosome clustering when individual lysosomes could still be resolved. This presumption was borne out by the extensive co-localization of spike and Lamp2 in small lysosomes scattered throughout the cytoplasm of control Htet1/S^D614G^ cells (***Fig. 4D***). As for whether spike-containing lysosomes were able to accumulate A647-dextran uptake similarly to other lysosomes in the cell, or in other cells, we observed that spike-containing lysosomes displayed little if any A647-dextran fluorescence, even in cells that contained many spike-negative, A647-dextran-positive lysosomes (***Fig. 4E, F***). Interestingly, both populations of lysosomes within these cells (spike+, A647- and spike-, A647+) appeared to cluster together in cells that had larger lysosome clusters.

Lysosome acidification is another hallmark of lysosome function, and is catalyzed by the vacuolar ATPase (V-ATPase) proton pump (Futai et al., 2019). A number of probes have been developed that label acidic compartments (Chazotte, 2011), including the lysotracker series of probes. To determine whether spike-containing lysosomes have a pH similar to spike-negative lysosomes, we labeled Htet1/S^D614G^ cells with Lysotracker Deep Red and then processed the cells for immunofluorescence microscopy using plasma G4 to detect the lysosomal forms of spike. Lysotracker Deep Red labeled many lysosomes brightly, and while some spike-containing lysosomes displayed labeling consistent with an acidic lumen, many of the spike-containing lysosomes displayed relatively low or no lysotracker fluorescences (***Fig. 4G***).

### Spike trafficking to lysosomes is resistant to inhibitors of endocytosis, microtubules, and secretion but sensitive to V-ATPase inhibition

Proteins can be trafficked to lysosomes following endocytosis from the plasma membrane by a pathway that often effects their destruction, or by direct intracellular vesicle traffic (Braulke and Bonifacino, 2009). To determine whether the lysosomal trafficking of spike occurs via the endocytic pathway so often associated with cargo protein destruction, we incubated Htet1/S^D614G^ cells with doxycycline for 14 hours in the presence of vehicle alone (DMSO) or in media containing either of two inhibitors of endocytosis, the dynamin-1 inhibitor dynasore, which blocks clathrin-dependent endocytosis (Kirchhausen et al., 2008), or the clathrin inhibitor pitstop2, which blocks both clathrin-dependent and clathrin-independent endocytosis (Dutta et al., 2012). Htet1/S^D614G^ cells incubated with vehicle alone (DMSO) localized spike to Lamp2-positive lysosomes (***Fig. 5A***) and the same was observed for Htet1/S^D614G^ cells incubated with either dynasore or pitstop2 (***Fig. 5B, C***).

**Figure 5.**
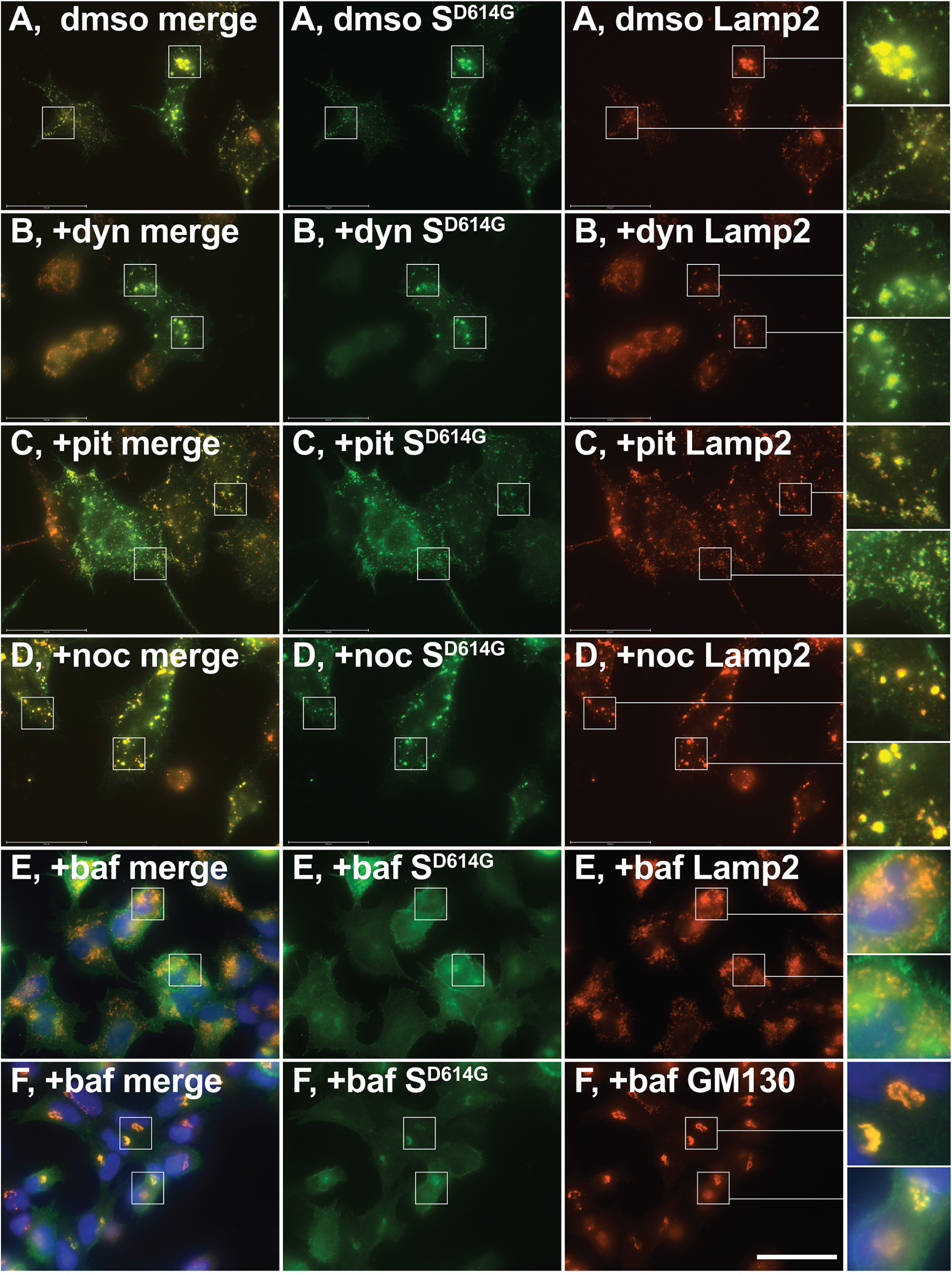
Spike expression induces lysosome clustering and inhibits lysosome function. (A-C) Micrographs of Htet1/S^D614G^ cells induced with doxycycline for two days followed by treatment with (A) DMSO or (B, C) vacuolin-1 for 3 hours. Cells were stained with (green) COVID-19 patient plasma G4, (red) a monoclonal anti-Lamp2 antibody, and (blue) DAPI. (D-F) Micrographs of Htet1/S^D614G^ cells induced with doxycycline for 20 hours (D) without treatment, (E) incubated with Alexa Fluor 647-dextran for an additional 3 hours in media containing doxycycline but lacking Alexa Fluor 647-dextran, or (F) incubated for one hour in media containing both doxycycline and Lysotracker Deep Red. Cells were stained with (green) COVID-19 patient plasma G4 and (red) a monoclonal anti-Lamp2 antibody. Insets (∼3-fold higher magnification) show greater detail in areas of particular interest. Bar, 50 μm.

The microtubule-inhibitor nocodazole, while not a general inhibitor of endocytosis, has been shown to impair the lysosomal uptake of selected endocytosed proteins (Tacheva-Grigorova et al., 2013). Moreover, it is known to affect lysosome function by blocking microtubule-dependent retrograde and anterograde movements of lysosomes (Cabukusta and Neefjes, 2018). Addition of nocodazole had no discernable effect on the lysosomal sorting of spike (***Fig. 5D***) and while it had some effect on lysosome distribution, it did not prevent spike-induced lysosome clustering. Lysosome function is more severely inhibited by bafilomycin A1, a potent inhibitor of the V-ATPase that blocks acidification of lysosomes (and other organelles), prevents the maturation of many lysosomal hydrolases, and inhibits the activity of many pH-dependent lysosomal enzymes. and required for lysosome acidification (Braulke and Bonifacino, 2009; Futai et al., 2019). To explore its effects on the lysosomal sorting of spike, cells were incubated with bafilomycin throughout the period of spike induction and then processed for immunofluorescence microscopy. Bafilomycin-treated cells localized spike to Golgi and plasma membranes rather than lysosomes (***Fig. 5F-I***), demonstrating that the trafficking of spike to lysosomes is a V-ATPase-dependent process. Interestingly, the lysosomes of bafilomycin-treated cells showed little if any clustering.

### The D614G mutation enhances the lysosomal sorting of spike

Given that the D614G mutation has a significant impact on SARS-CoV-2 infectivity and transmission, we asked whether this mutation might impact the lysosomal sorting of spike. Htet1/S^W1^ and Htet1/S^D614G^ cells express spike in response to doxycycline, and at similar levels (***Fig. 6A***). Significant amounts of spike are proteolytically processed at its 682RRAR685 motif to generate the S2 and S1 fragments, and these cells also contained similar levels of both S2 and S1. However, the D614G mutation did induce a slight shift in the relative extent of processing at the S1/S2 boundary, shown here by somewhat less full-length spike in Htet1/S^D614G^ cells relative to Htet1/S^W1^ cells.

**Figure 6.**
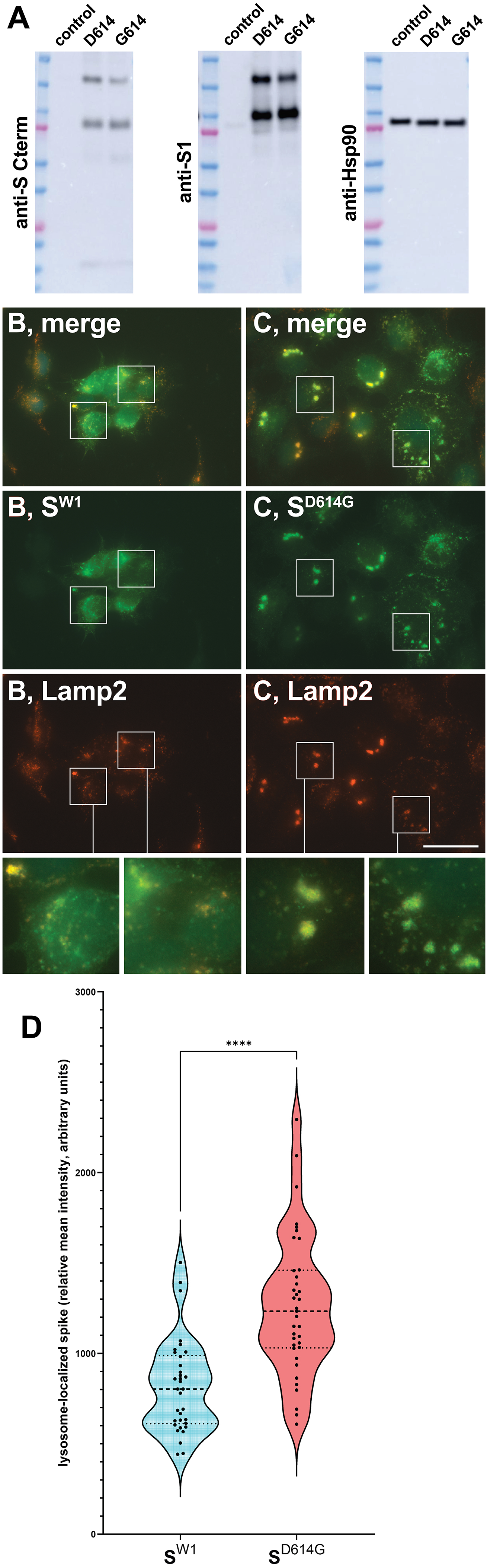
The D614G mutation enhances lysosomal sorting of spike. (A) Immunoblot analysis of doxycycline-treated Htet1, Htet1/S^W1^, and Htet1/S^D614G^ cells. Cell lysates were prepared and separated by SDS-PAGE, then processed for immunoblot using (left panel) affinity purified antibodies specific for the C-terminal 14 amino acids of spike, (center panel) a mouse monoclonal antibody specific directed against the S1 region of spike, and (right panel) endogenously expressed Hsp90. (B, C) Fluorescence micrographs of doxycycline-treated Htet1/S^W1^ and Htet1/S^D614G^ cells stained with (green) affinity purified antibodies specific for the C-terminal 14 amino acids of spike, (red) a mouse monoclonal antibody specific for Lamp2, and (blue) DAPI. The upper panels (merge) display the composite of all three images, above the individual images for (green) spike and (red) Lamp2. Insets (∼3-fold higher magnification) show greater detail in areas of particular interest. Bar, 50 μm. (D) Plot of lysosome-associated levels of spike (relative mean fluorescence, arbitrary units) in cells expressing S^W1^ or S^D614G^.

To determine the subcellular distribution of the D614 and G614 forms of spike, doxycycline-induced Htet1/S^W1^ and Htet1/S^D614G^ cells were processed for immunofluorescence microscopy using an antibody that binds the C-terminal 14 amino acids of spike (-DSEPVLKGVKLHYT_COOH_), a region of the protein that is shielded by a lipid bilayer from the D614G mutation and the conformational changes it might induce. Although both forms of spike displayed significant co-localization with the lysosomal marker Lamp2, the D614G mutation appeared to induce a shift in spike protein sorting towards the lysosome (***Fig. 6B, C***). Although the relative amounts of lysosome-localized spike varied significantly from one cell to another in both cell populations, digital image analysis quantified the effect of the D614G mutation as 54% increase in its lysosomal staining (*p* = 0.00000024; Student’s *t-*test; 2-tailed; two-sample, unequal variance), from an average of 817 +/− 44 (standard error of the mean (s.e.m.)) in Htet1/S^W1^ cells (n = 34 images) to an average of 1265 +/− 64 (s.e.m) in Htet1/S^D614G^ cells (n = 37 images) (***Fig. 6D***).

These results suggest a model of SARS-CoV-2 biogenesis in which the D614G mutation enhances spike protein trafficking to the lysosome. Given the recent report that the biogenesis of newly-synthesized MHV particles occurs via lysosomes and that MHV-infected cells accumulate MHV particles in lysosomes (Ghosh et al., 2020), we tested whether the D614G mutation might promote the lysosomal accumulation of spike and spike-containing vesicles in SARS-CoV-2-infected cells. More specifically, we infected VeroE6TMPRSS2 cells with equal m.o.i. of a G614 strain of SARS-CoV-2 (HP7) or a D614 strain of SARS-CoV-2 (HP76). 18 hours after initiation of infection, the two cell populations were fixed, permeabilized, and processed for confocal immunofluorescence microscopy using antibodies specific for spike and Lamp2 (***Fig. 7***). In cells infected with the G614 virus, spike was localized primarily to small intracellular vesicles clustered in the perinuclear area of the cell, proximal to Lamp2-positive structures, often co-localizing with Lamp2, and especially in small, virus-sized punctae surrounded by Lamp2-positive membrane (***Fig. 7A-D***). In contrast, spike distribution in D614 virus-infected cells was distributed across a much broader array of intracellular compartments, especially the plasma membrane and intracellular compartments distant from Lamp2-positive structures (***Fig. 7E-H***). However, these differences were a matter of degree, as D614 spike also displayed some labeling in Lamp2-positive and nearby compartments, as well as in small spike-positive puncta surrounded by Lamp2-containing membrane.

**Figure 7.**
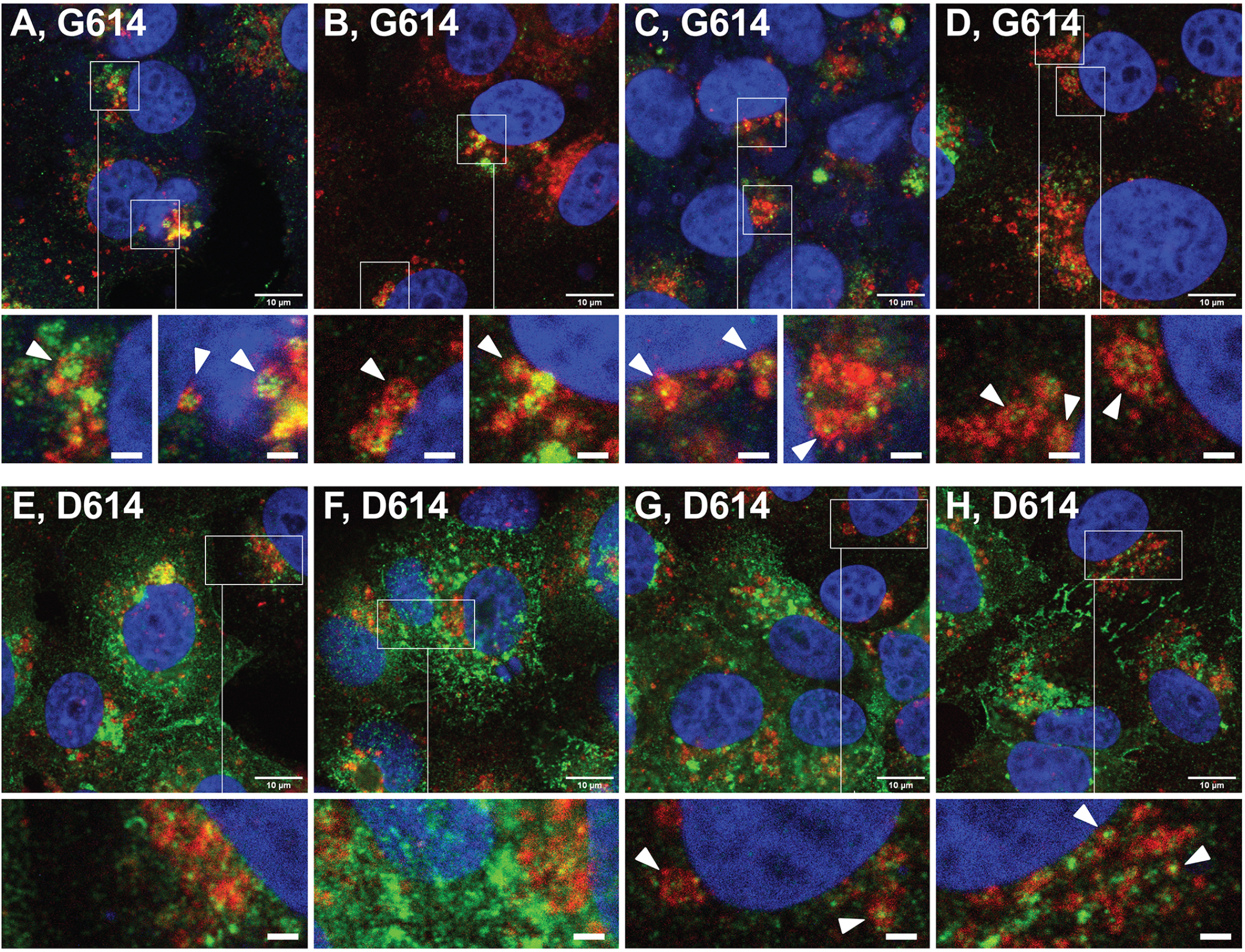
G614 and D614 spike in SARS-CoV-2-infected cells. Confocal fluorescence micrographs of Vero/TMPRSS2 cells infected with (A-D) a spike G614 strain of SARS-CoV-2 or (E-H) a spike D614 strain of SARS-CoV-2. Cells were infected with equal amounts of virus, incubated for 18 hours, then fixed, permeabilized, and stained with (green) affinity purified antibodies specific for the C-terminal 14 amino acids of spike, (red) a monoclonal anti-Lamp2 antibody, and (blue) DAPI. Insets (∼3-fold higher magnification) show greater detail in areas of particular interest. Bar in original images, 10 μm; Bar in magnified insets, 2 μm.

## Discussion

We show here that the spike D614G mutation alters spike protein sorting, enhancing its trafficking towards the lysosome and away from other intracellular compartments. This D614G-induced shift in spike protein trafficking was observed in human cells expressing the spike protein on its own, outside the context of a viral infection, where it appeared to enhance spike-induced lysosomes clustering. It was also observed in the context of SARS-CoV-2-infected cells, as spike protein encoded by G614 virus displayed increased association with lysosomes and reduced accumulation in non-lysosomal compartmets and areas of the cell. Furthermore, we observed that cells infected by a G614 strain of SARS-CoV-2 exhibited an enhanced accumulation of discrete, small, spike-containing punctae within Lamp2-positive membranes. In addition, we observed that the D614G mutation induced a slight elevation in spike protein processing at the S1/S2 junction. To the best of our knowledge, the D614G-induced lysosomal shift in spike protein trafficking and slight elevation in spike processing represent the earliest cell biological and biochemical manifestations of the D614G mutation, raising the strong possibility that they may contribute to the pronounced effects of the D614G mutation on SARS-CoV-2 infectivity, viral load, and transmission (Hou et al., 2020; Korber et al., 2020; Lorenzo-Redondo et al., 2020).

Numerous lines of evidence support our conclusion that spike is trafficked to the lysosome membrane, including its co-localization with Lamp1, Lamp2, and Lamp3/CD63, its ring-like staining in vacuolin-1-treated cells, its induction of lysosome clustering, and its lysosomal localization when detected with a relatively conformation-independent antibody probe (the affinity purified antibodies to its C-terminal 14 amino acids. As for how spike is trafficked to lysosomes, our data indicate that it can occur independently of endocytosis, as it’s lysosomal accumulation was not blocked by inhibitors of clathrin-dependent or clathrin-independent endocytosis. Spike trafficking to lysosomes was also unaffected by the microtubule depolymerizing agent nocodazole, even though nocodazole ablates motor-directed lysosome movements. Spike trafficking to lysosomes was, however, strongly inhibited by bafilomycin A1, a V-ATPase inhibitor, indicating that compartment acidification is critical to the lysosomal trafficking of spike. These results are consistent with those from a recent CRISPR-based screen for genes required for SARS-CoV-2 replication, which identified 13 components of the V-ATPase among the 50-most important genes for SARS-CoV-2 replication (Daniloski et al., 2020).

In addition to being trafficked to lysosomes, the expression of spike appears to alter lysosome function. This was apparent from the fact that spike expression induces a pronounced clustering of lysosomes, especially in cells expressing the G614 form of spike (***Table 2***). While we do not know the molecular mechanism by which spike expression drives lysosome clustering, it appeared to be a microtubule-independent process, as it was not inhibited by nocodazole, similar to previous studies of lysosome clustering (Ba et al., 2018). Spike-induced alteration of lysosome function was also apparent from was the fluorescent dextran uptake experiments, which revealed that spike-containing lysosomes were defective in uptake of this fluid-phase endocytosis marker, and may also display a somewhat reduced ability to acidify their lumen, as some spike-containing lysosomes displayed low labeling with Lysotracker Deep Red.

**Table 2.**
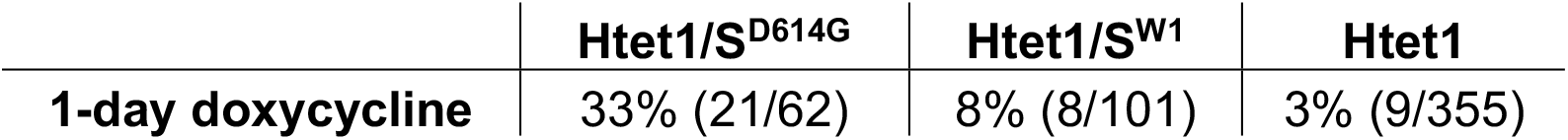
Percentage of cells with clustered (Lamp2+) lysosomes.

Given that the D614G mutation enhances viral fitness, it is important to consider how viral fitness might be promoted by an increase in the lysosomal sorting of spike and the accumulation of spike-containing punctae within lysosomes of SARS-CoV-2-infected cells. The most obvious possibility is that these changes facilitate the lysosomal phases of SARS-CoV-2 virion biogenesis. This conclusion is based in part on the recent report by Ghosh et al that MHV is released independently of the biosynthetic secretory pathway, that MHV virions accumulate in lysosomes, that MHV release is an Arl8-dependent lysosomal secretory pathway, and that SARSW-CoV-2 virions accumulate in a lysosome-like structure of SARS-CoV-2-infected cells (Ghosh et al., 2020). This model of coronavirus egress involves some degree of aberration to protein egress in general, as MHV infection induced the lysosomal trafficking of the KDEL receptor (Ghosh et al., 2020), a phenotype that we observed for spike-expressing cells.

An analysis of matched D614 and G614 SARS-CoV-2 virions revealed no difference in the ultrastructure of SARS-CoV-2 particles, the shape of spike trimers, or the amounts of S and S2 in viral particles (Hou et al., 2020), while Zhang et al. (doi: 10.1101/2020.06.12.148726) concluded that the D614G mutation led to enhanced retention of S1. Our data show that cells expressing matched D614 and G614 forms of spike show no difference in the amounts of cell-associated S1 and S2, though there may be a slightly lower amount of full-length S. This may reflect a slight increase in the cleavage of G614 spike at the S1/S2 boundary, and since lysosomal proteases can catalyze this event it may be a consequence of enhanced lysosomal sorting of D614G spike proteins. Alternatively, enhanced spike cleavage by Golgi proteases such as furin may potentiate the lysosomal sorting of spike.

The most obvious effect of the D614G mutation is the ∼4-8-fold increase in a proxy marker of SARS-CoV-2 infection (an encoded luciferase) at 8 hours post-infection (Hou et al., 2020), suggestive of accelerated viral entry. Although it is at least formally possible that these various effect of the D614G mutation are unrelated, the more parsimonious hypothesis is that they are all reflections of a common mechanism. As for what this mechanism might be, there is already extensive precedent from the field of lysosomal protein sorting. Specifically, the mannose-6-phosphate (M6P)-M6P receptor system mediates the lysosomal delivery of both newly-synthesized intracellular cargoes and endocytosed cargoes retrieved from the extracellular milieu (Braulke and Bonifacino, 2009). Although there is a report of M6P modification on spike glycans (Brun et al. 2020 https://www.biorxiv.org/content/10.1101/2020.11.16.384594v1.full.pdf), we do not suggest that the effects of the D614G mutation are necessarily mediated by M6P receptors. Rather, we posit the more general hypothesis that spike protein interactions with one or more cell surface proteins accelerate SARS-CoV-2 entry and that the same or similar interactions facilitate the lysosomal trafficking of newly-synthesized spike proteins, perhaps by interactions with newly-synthesized versions of the same spike-binding proteins in the ER, ERGIC, or Golgi. In light of this hypothesis, it is interesting to note that SARS-CoV-2 infection is exquisitely sensitive to a known inhibitor of plasma membrane-to-lysosome traffic (Kang et al., 2020) and that SARS-CoV-2 infection requires numerous proteins involved in lysosome biology (Daniloski et al., 2020).

## Materials and Methods

### Cell lines, cell culture, transfections

HEK293 cells (ATCC) were cultured in complete medium (DMEM containing 10% fetal bovine serum and 1% penicillin/streptomycin solution (10,000 units/ml)). Transfections were carried out using lipofectamine according to the manufacturer’s instructions. Htet1 cells were generated by transfecting HEK293 cells with the plasmid pS147, which encodes the tetracycline-activated transcription factor rtTAv16 (Zhou et al., 2006), followed by selection of zeocin-resistant transgenic cell clones (200 ug/ml zeocin), and pooling of these clones. SARS-CoV-2 protein-expressing cell lines were generated by transfection of Htet1 cells with Sleeping Beauty transposons carrying a tet-regulated transgene designed to express each SARS-CoV-2 protein under control of the doxycycline-regulated TRE3G promoter. One to two days after transfection, cells were placed in selective media containing 1 ug/ml puromycin and 200 ug/ml zeocin. Multiple clones were obtained from each transfection, pooled, and expanded to create master banks of each test line. Expression of SARS-CoV-2 protein was induced by adding doxycycline to the culture medium at a final concentration of 1 ug/ml.

### Plasmids

The plasmid pS147 is a CMV-based vector designed to express the rtTAv16 protein from a polycistronic ORF, upstream of a viral 2a peptide and the Bleomycin resistance coding region. Other vectors used in this study were based on a Sleeping Beauty transposon vector (pITRSB) in which genes of interest can be inserted between the left and right inverted tandem repeats (ITRs). These include three plasmids in which the region between the ITRs contains (a) one gene in which a crippled EF1alpha promoter drives expression of a polycistronic ORF encoding mCherry, the p2a peptide, and the puromycin-resistance protein, and (b) a second gene in which the TRE3G promoter drives expression of codon-optimized forms of the N, S**, or M proteins (pCG217, pCG218, and pCG221, respectively, which were used to create the Htet1/N, Htet1/S**, and Htet1/M cell lines). Two additional transposon-mobilizing plasmids were also used in this study. These plasmids carry (a) one gene in which a crippled EF1alpha promoter drives expression of the puromycin-resistance protein, and (b) a second gene in which the TRE3G promoter drives expression of codon-optimized forms of the S^W1^ or S^D614G^ proteins (pCG145 and pCG200, respectively, used to create the Htet1/ S^W1^ and Htet1/ S^D614G^ cell lines). All plasmid sequences are available upon request. S** encodes the same protein as S^W1^, with the exception of 6 amino acid changes (986KV987 to 986PP987 and 682RRAR685 to 682GSAG685). S^D614G^ encodes the same protein as S^W1^, with the exception of the D614G amino acid substitution.

### Immunoblot

HEK293 cell lines were grown in the presence or absence of doxycycline, lysed by addition of sample buffer, separated by SDS-PAGE, transferred to PVDF membranes, and incubated with primary antibodies and HRP-conjugated secondary antibodies. Following extensive washes, proteins were membranes visualized using chemiluminescence reagents and an Amersham Imager 600 gel imaging system.

### Immunofluorescence microscopy

Cells were cultured on either sterile, poly-L-lysine-coated coverglasses, or sterile, poly-L-lysine-coated, glass-bottom, black-walled 96 well plates. For serology testing, SARS-CoV-2 protein-expressing cells were mixed with the parental Htet1 cell line at a ratio of ∼90%:10%. Cells were exposed to 1 ug/ml doxycycline for 1 day to induce SARS-CoV-2 protein expression. Cells were then fixed (4% formaldehyde in PBS), permeabilized (1% Triton X-100 in PBS), and processed for immunofluorescence microscopy using established protocols. Coverglasses were mounted on slides using Fluoromount G (catalog #17984-25, Electron Microscopy Sciences). Stained cells were visualized using an EVOSM7000 fluorescence microscope (ThermoFisher) equipped with 20x (PL FL 20X, 0.50NA/2.5WD), 40x (PLAN S-APO 40X, NA0.95, 0.18MM), and 60x (OBJ PL APO 60X, 1.42NA/0.15WD) Olympus objectives) Confocal fluorescence microgaphs were acquired using a Zeiss LSM800 microscope with gallium-arsenide phosphide (GaAsP) detectors and a 100x/1.4na Plan-Apochromat objective. Images were assembled into figures using Adobe Illustrator. Quantitative analysis of image files was performed using ImageJ and proprietary software.

### Antibodies and other reagents

Rabbit polyclonal antibodies were raised against synthetic peptides corresponding to carboxy-terminal peptides of SARS-CoV-2 N (-KQLQQSMSSADSTQA_COOH_) and S (-DSEPVLKGVKLHYT_COOH_). Rabbit antibodies to the SARS M protein carboxy-terminal peptide (-DHAGSNDNIALLVQ_COOH_) were a gift from C. Machamer, Johns Hopkins University. ThermoFisher was the source for rabbit antibodies directed against LAMTOR1 (#8975S), ERGIC3 (#16029-1-AP), ERGIC53 (#13364-1-AP), and calnexin (#PA5-34665) and for mouse monoclonal antibodies to EEA1 (#48453) and Lamp2 (MA1-205). Rabbit antibodies to mTOR were obtained from Cell Signaling (#2972S). Mouse monoclonal antibodies to GM130 (#610822) and CD81 (#555675) were obtained from BD. Mouse monoclonal antibody to Lamp3/CD63 (#NBP2-32830) was obtained from Novus. Mouse monoclonal antibody directed against CD9 (#312102) was obtained from BioLegend. Rabbit polyclonal antibodies to BiP/GRP78 (#21685) and mouse monoclonal to Lamp1 (#24170) were from Abcam. Fluorescently labeled secondary antibodies specific for human Igs (IgG, IgM, and IgA; pan Ig), human IgG, rabbit IgG, or mouse IgG were obtained from Jackson ImmunoResearch. Lysotracker Deep Red (#L12492) and Alexa647-Dextran 10,000 MW (#D22914) were obtained from ThermoFisher. We thank Dr. Nihal Altan-Bonnet for generous gift of the anti-BiP/GRP78 antibodies, the Lamp1 monoclonal, and polyclonal antibody to the KDEL receptor.

### Human Plasmas

All patient plasmas were collected using standard procedures for blood draw and plasma collection. Following Johns Hopkins Medicine Institutional Review Board (IRB) approval, plasma samples were obtained under informed consent from healthy donors prior to the COVID-19 pandemic (JHM IRB NA_0004638) as described (Cox et al., 2005). The COVID-19 specimens utilized for this publication were obtained from the Johns Hopkins Biospecimen Repository, which is based on the contribution of many patients, research teams, and clinicians, and were collected following IRB approval (Johns Hopkins COVID-19 Clinical Characterization Protocol for Severe Infectious Diseases (IRB00245545) and Johns Hopkins COVID-19 Remnant Specimen Repository (IRB00248332)). All COVID-19 patient plasmas used in this study were collected on the day of admission of the patient into the Johns Hopkins Hospital, and between the dates of April 7 and April 22, 2020.

### Virus and virus infections

VeroE6TMPRSS2 cells (Matsuyama et al., 2020) were used to grow and titrate infectious virus using established protocols (Klein et al., 2020; Schaecher et al., 2007). The clinical isolates SARS-CoV-2/USA/MD-HP00076/2020 (Spike D614; GenBank: MT509475.1) and SARS-Cov-2/USA/DC-HP00007/2020 (Spike G614; GenBank: MT509464.1) were isolated using published procedures (Gniazdowski et al., 2020) and virus stocks were grown on VeroE6TMPRSS2 cells. Virus stocks were sequenced to confirm that the amino acid sequence of the isolate was identical to the sequence derived from the clinical sample.

### Sources of Support

Funding for this work was provided by Johns Hopkins University, by Capricor, and by grants from the National Institutes of Health (NIH) National Cancer Institute (UG3CA241687), the NIH Center of Excellence in Influenza Research and Surveillance (HHSN272201400007C), and the National Institute of Allergy and Infectious Diseases (U19A1088791).

## Acknowledgments

The authors thank Dr. Carolyn Machamer and Nihal Altan-Bonnet for their numerous insights and suggestions during the course of this work, as well as for their kind gift of antibodies. The authors acknowledge Dr. LaToya A. Roker (Johns Hopkins School of Medicine Microscope Facility) for excellent assistance with confocal image acquisition and processing, James Morrell for expert assistance with plasmid assembly, and Guido Massaccesi for assistance in plasma sample handling. We thank the National Institute of Infectious Diseases, Japan, for providing VeroE6TMPRSS2 cells. The specimens utilized for this publication were part of the Johns Hopkins Biospecimen Repository, which is based on the contribution of many patients, research teams, and clinicians.

## Legends to Supplemental Figures

**Supplemental figure 1.**
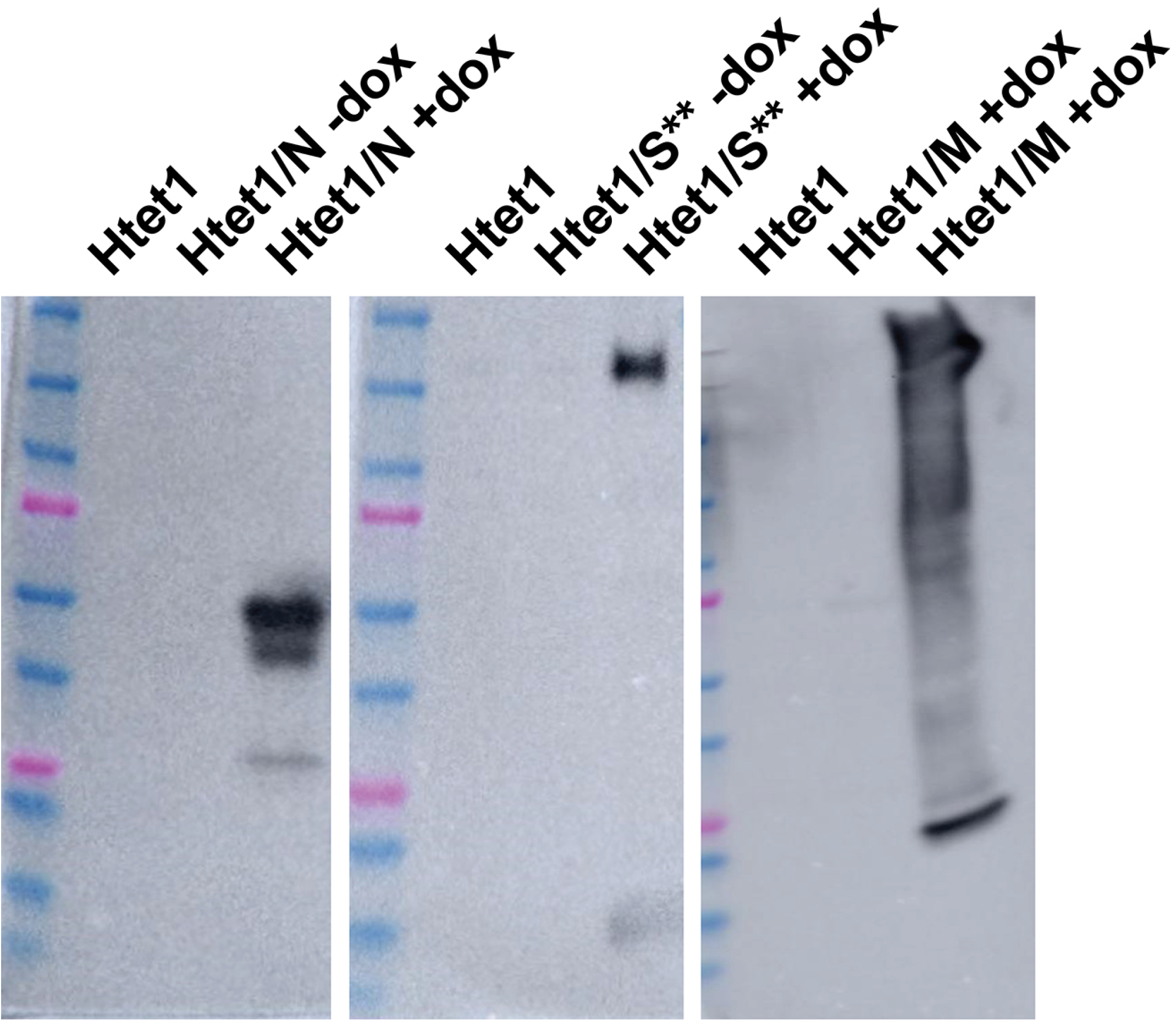
Doxycycline-inducible expression of SARS-CoV-2 N, S**, and M. Immunoblot of whole cell lysates of Htet1 cells, and of Htet1/N, Htet1/S**, and Htet1/M cells grown in the absence or presence of doxycycline. Blots were probed with (left panel) rabbit polyclonal anti-peptide antibodies specific for SARS-CoV-2 N protein, (center panel) rabbit polyclonal anti-peptide antibodies specific for SARS-CoV-2 S spike protein, and (right panel) rabbit polyclonal anti-peptide antibodies specific for the SARS and SARS-CoV-2 M protein. Size markers, from top: 250 kDa, 150 kDa, 100 kDa, 75 kDa, 50 kDa, 37 kDa, 25 kDa, 20 kDa, 15 kDa, 10 kDa. Predicted molecular masses are 46 kDa for N, 141 kDa for S**, and 25 kDa for M. The high MW forms of M apparent in these blots were observed whenever M was expressed in isolation.

**Supplemental figure 2.**
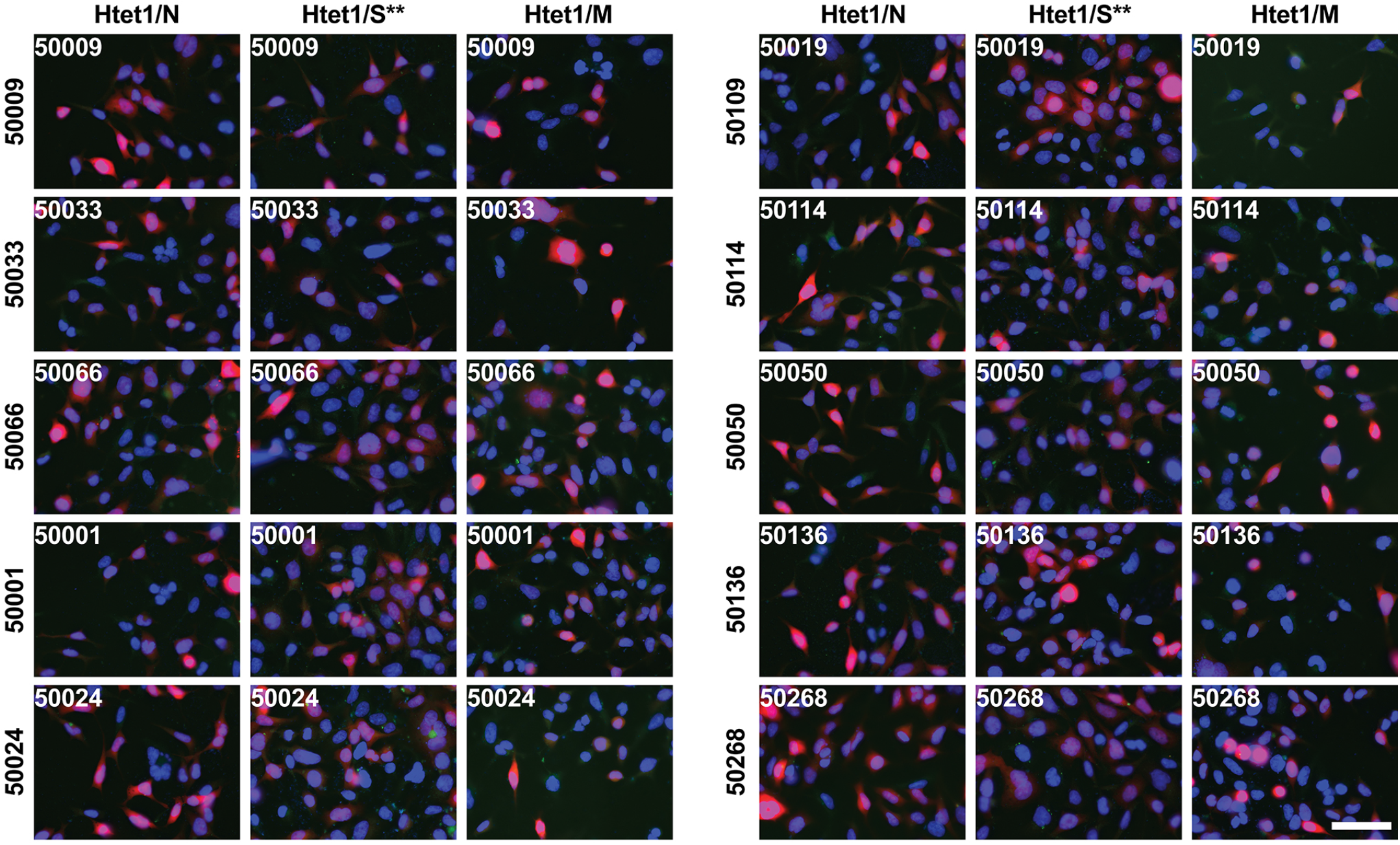
Control plasma staining data I. Fluorescence micrographs of doxycycline-induced Htet1/N, Htet1/S**, and Htet1/M cells. Cells were processed for immunofluorescence microscopy and stained to detect (green) bound human plasma Ig and (blue) DAPI, then imaged to detect bound Ig, (red) mCherry, and DAPI. Micrographs present the merge of all three labels and are organized with cell lines in each column and plasmas in each row. These plasmas were collected prior to COVID-19 pandemic. Bar, 75 μm.

**Supplemental figure 3.**
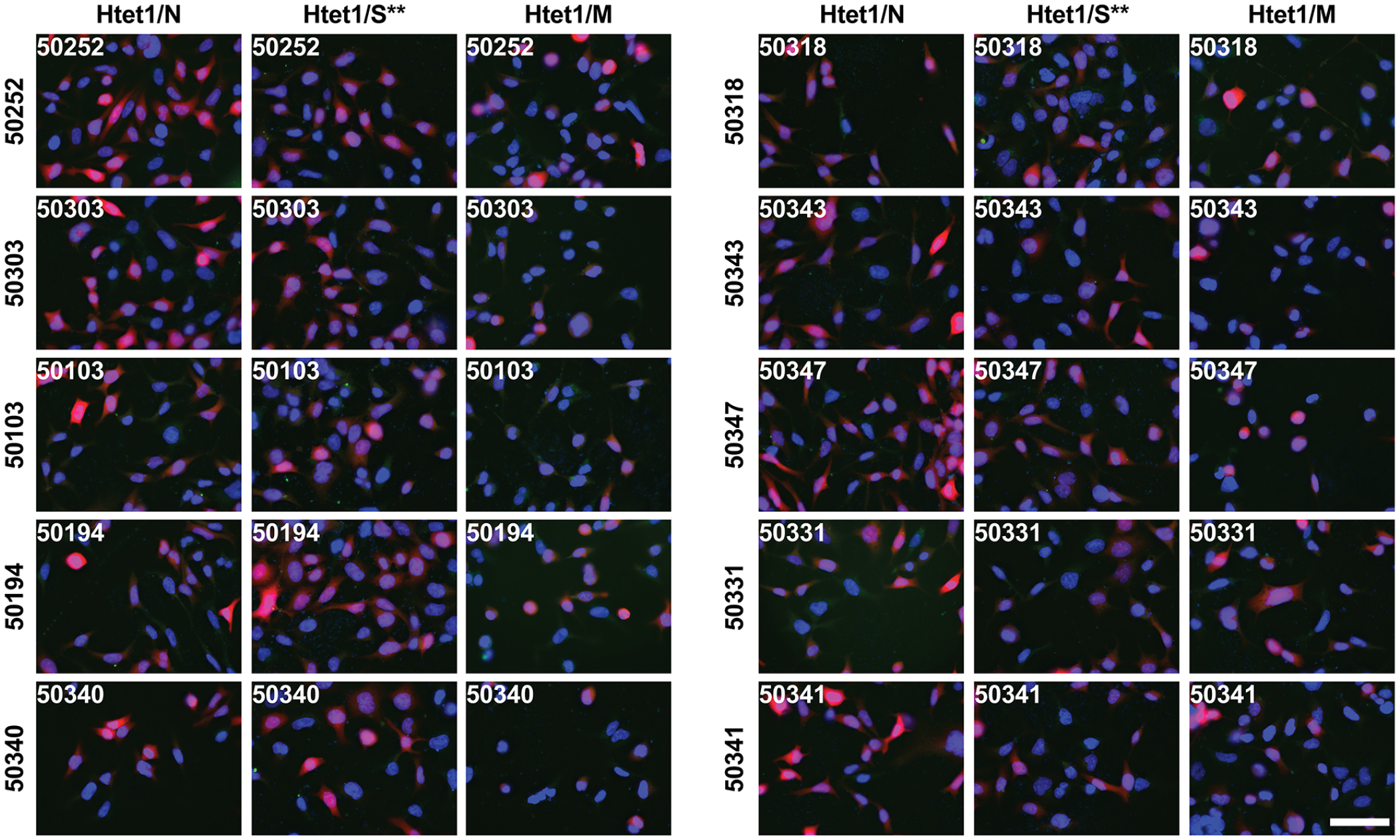
Control plasma staining data II. Fluorescence micrographs of doxycycline-induced Htet1/N, Htet1/S**, and Htet1/M cells. Cells were processed for immunofluorescence microscopy and stained to detect (green) bound human plasma Ig and (blue) DAPI, then imaged to detect bound Ig, (red) mCherry, and DAPI. Micrographs present the merge of all three labels and are organized with cell lines in each column and plasmas in each row. These plasmas were collected prior to COVID-19 pandemic. Bar, 75 μm.

**Supplemental figure 4.**
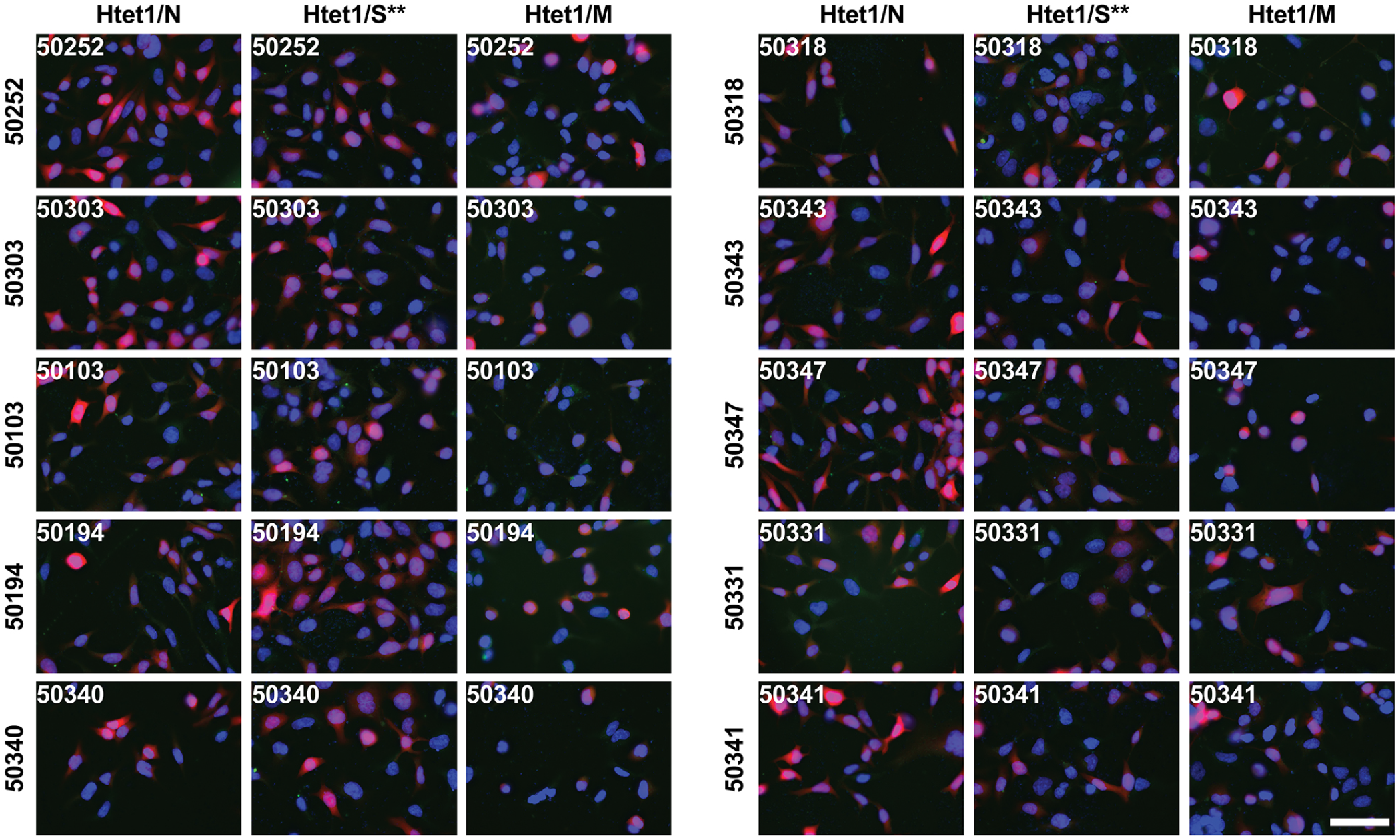
Control plasma staining data III. Fluorescence micrographs of doxycycline-induced Htet1/N, Htet1/S**, and Htet1/M cells. Cells were processed for immunofluorescence microscopy and stained to detect (green) bound human plasma Ig and (blue) DAPI, then imaged to detect bound Ig, (red) mCherry, and DAPI. Micrographs present the merge of all three labels and are organized with cell lines in each column and plasmas in each row. These plasmas were collected prior to COVID-19 pandemic. Bar, 75 μm.

**Supplemental figure 5.**
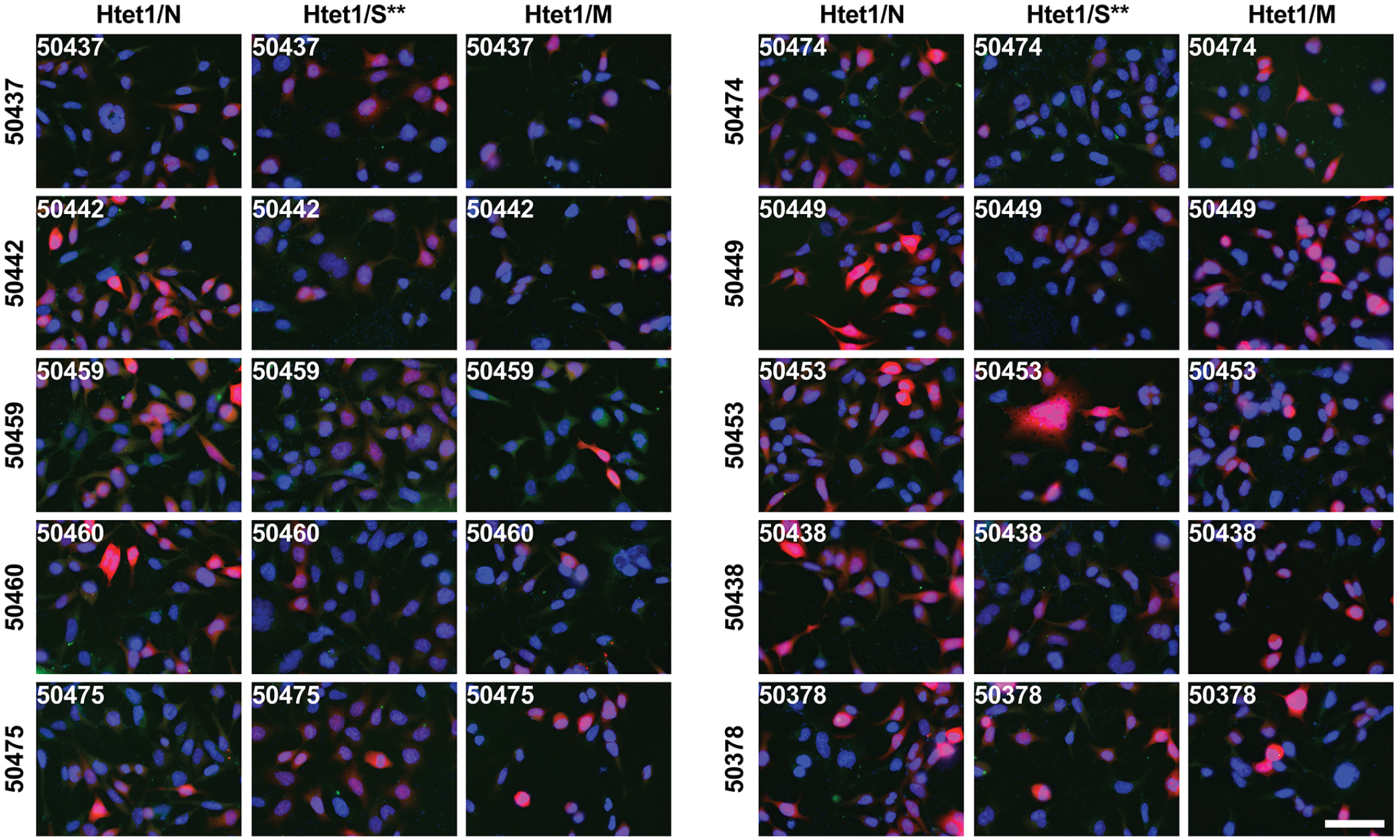
Control plasma staining data IV. Three-color merged fluorescence micrographs of Htet1/N, Htet1/S**, and Htet1/M cells imaged for (green) human plasma Ig staining, (red) mCherry expression and (blue) DAPI. Micrographs are organized with cell lines in each column, and plasmas in each row. These plasmas were collected prior to COVID-19 pandemic. Bar, 75 μm.

**Supplemental figure 6.**
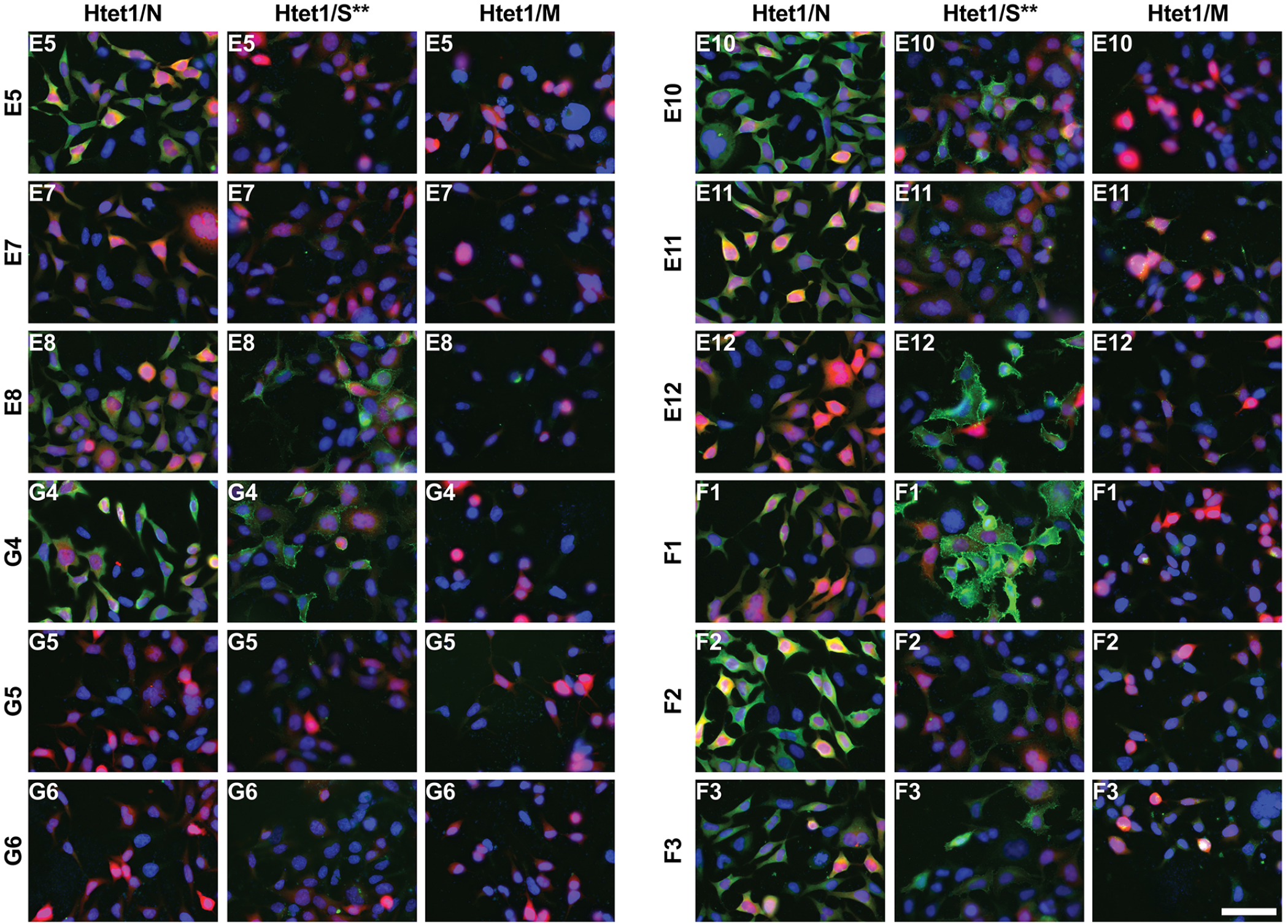
COVID-19 plasma staining data I. Three-color merged fluorescence micrographs of Htet1/N, Htet1/S**, and Htet1/M cells imaged for (green) bound human plasma Ig, (red) mCherry expression and (blue) DAPI. Micrographs are the merge of all three images and are organized with cell lines in each column, and plasmas in each row. These plasmas were collected from COVID-19 patients on their first day of admittance into Johns Hopkins Hospital in April 2020. Bar, 75 μm.

**Supplemental figure 7.**
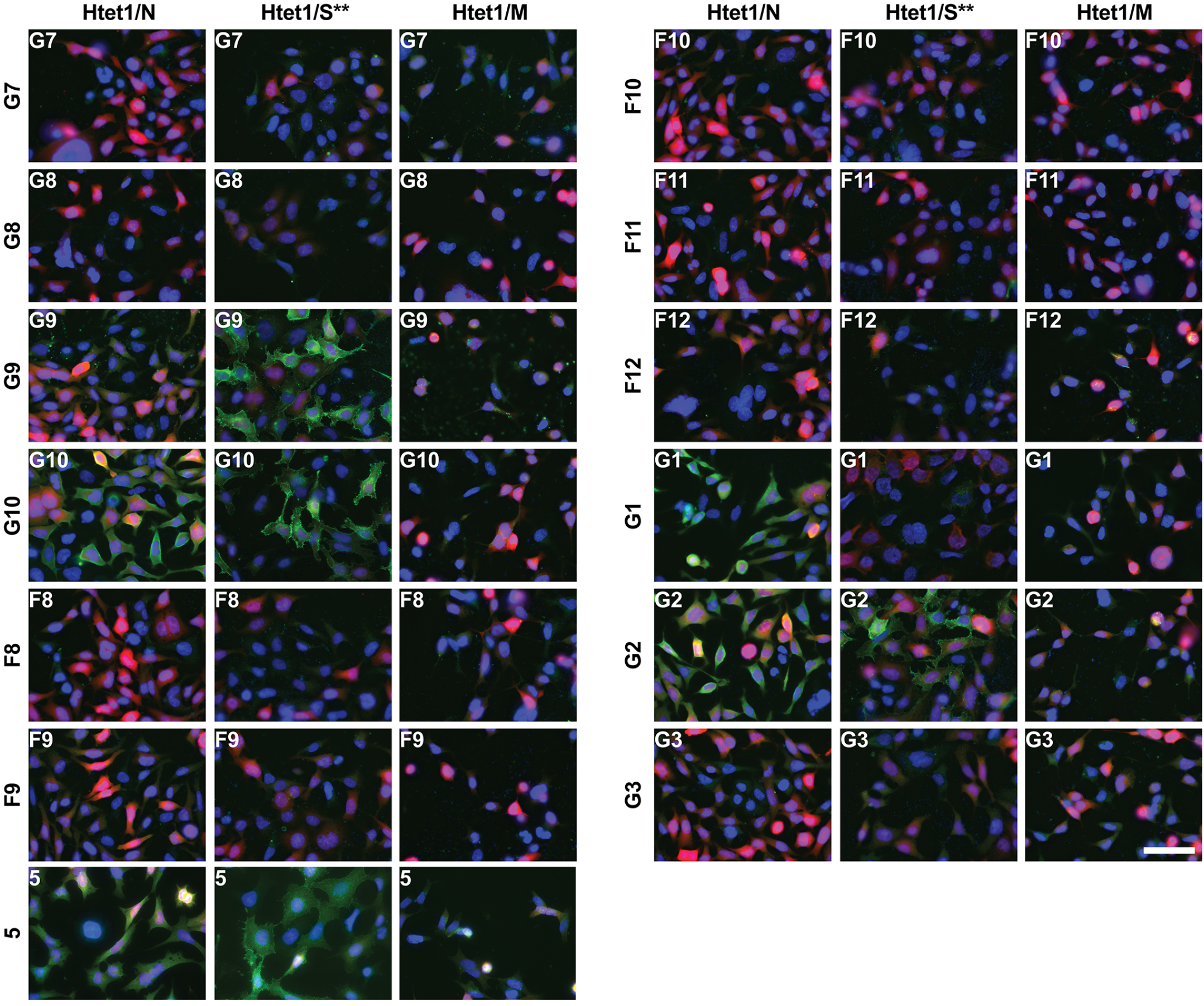
COVID-19 plasma staining data II. Three-color merged fluorescence micrographs of Htet1/N, Htet1/S**, and Htet1/M cells imaged for (green) bound human plasma Ig, (red) mCherry expression and (blue) DAPI. Micrographs are the merge of all three images and are organized with cell lines in each column, and plasmas in each row. These plasmas were collected from COVID-19 patients on their first day of admittance into Johns Hopkins Hospital in April 2020. Bar, 75 μm.

**Supplemental figure 8.**
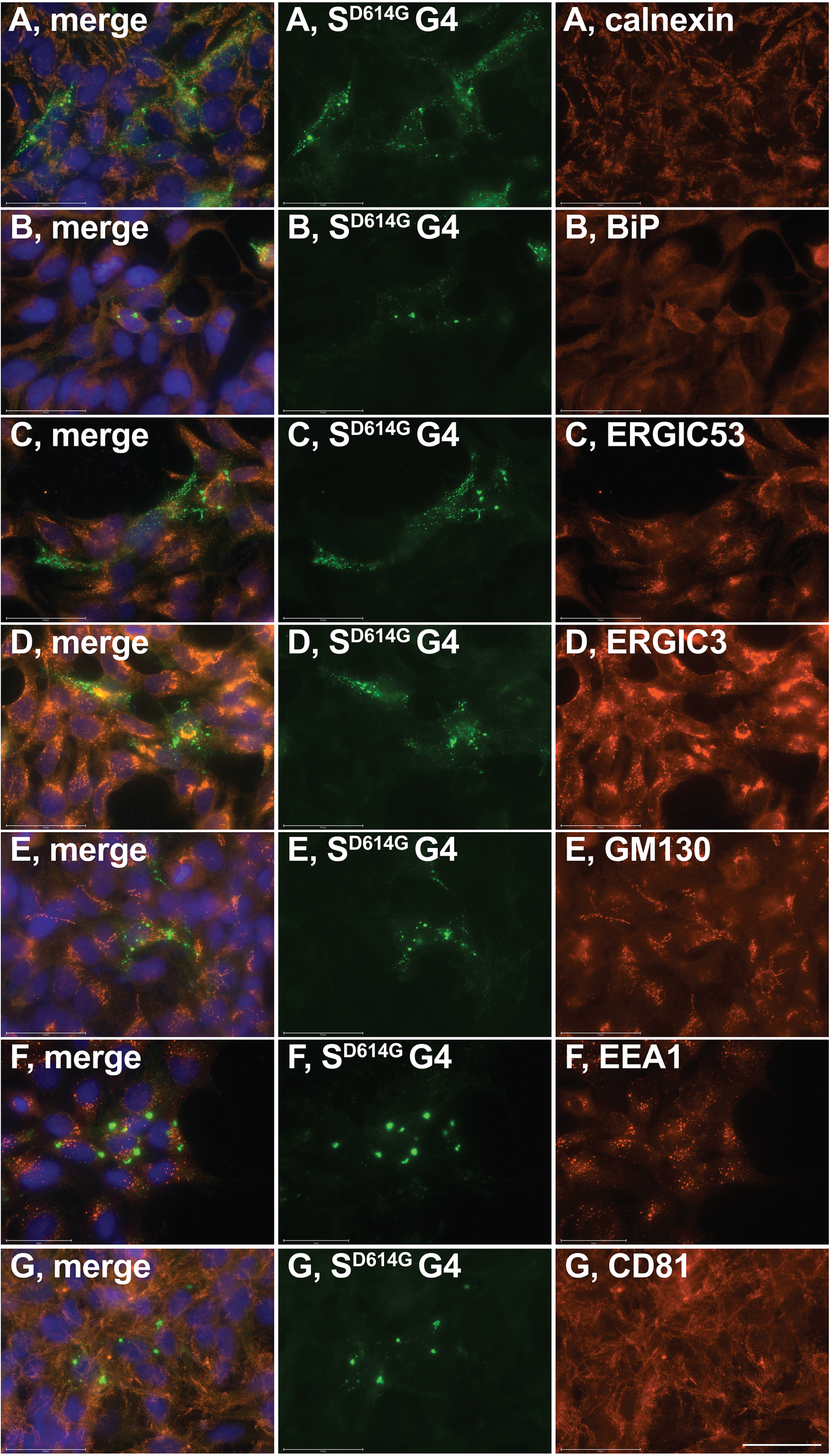
ER, ERGIC, Golgi, endosome, and plasma membrane marker proteins are not enriched in large, spike-containing intracellular compartments. Fluorescence micrographs of Htet1/S^D614G^ cells induced with doxycycline and processed for immunofluorescence microscopy using (blue) DAPI, (green) plasma G4, and (red) antibodies specific for (A) calnexin, (B) BiP/GRP78, (C) ERGIC53, (D) ERGIC3, (E) GM130, (F) EEA1, and (G) CD81. Bar, 50 μm.

